# Novel Peptide Inhibitor of Human Tumor Necrosis Factor-α has Antiarthritic Activity

**DOI:** 10.1101/2022.12.06.519274

**Authors:** Debasis Sahu, Charu Gupta, Ragothaman Yennamalli, Shikha Sharma, Saugata Roy, Sadaf Hasan, Pawan Gupta, Vishnu Kumar Sharma, Sujit Kashyap, Santosh Kumar, Ved Prakash Dwivedi, Amulya Kumar Panda, Hasi Rani Das, Chuan-Ju Liu

**Affiliations:** Product Development Cell, National Institute of Immunology, New Delhi, India; Department of Orthopedics Surgery, New York University School of Medicine, New York, NY, United States; Clinical Research Division, School of Basic and Applied Sciences, Galgotias University, Greater Noida, Uttar Pradesh, India; SASTRA Deemed to be University, Thanjavur, Tamil Nadu; Amity Institute of Forensic Sciences, Amity University, Noida, U.P, India; CSIR-Institute of Genomics and Integrative Biology, Delhi, India; Department of Pharmaceutical Chemistry, Shri Vile Parle Kelavani Mandal’s Institute of Pharmacy, Dhule, India; Dept of Pharmacoinformatics, National Institute of Pharmaceutical Education and Research (NIPER), Mohali, Punjab, India; Division of Pediatric Rheumatology, University of California San Francisco, CA, USA; Department of Genetics, University of Delhi, Delhi, India; Immunobiology Group, International Centre for Genetic Engineering and Biotechnology (ICGEB), New Delhi, India

**Keywords:** Tumor Necrosis Factor (TNF)-α, inflammation, peptide, nuclear factor kappa B (NFκB), collagen-induced arthritis (CIA)

## Abstract

The inhibition of tumor necrosis factor-α (TNFα) trimer formation renders it inactive for binding to its receptors thus mitigating the vicious cycle of inflammation. We designed a peptide (PIYLGGVFQ) that simulates a sequence strand of human TNFα monomer using a series of *in silico* methods, such as active site finding (Acsite), protein-protein interaction (PPI), docking studies (GOLD and Modeller) followed by molecular dynamics (MD) simulation studies. The MD studies confirmed the intermolecular interaction of the peptide with the TNFα. Fluorescence-activated cell sorting (FACS) and fluorescence microscopy revealed that the peptide effectively inhibited the binding of TNF to the cell surface receptors. The cell culture assays showed that the peptide significantly inhibited the TNFα-mediated cell death. In addition, the nuclear translocation of the nuclear factor kappa B (NFκB) was significantly suppressed in the peptide-treated A549 cells as observed in immunofluorescence and gel mobility-shift assays. Furthermore, peptide protected against joint damage in collagen-induced arthritis (CIA) mouse model as revealed in the microfocal-CT scans. In conclusion, this TNFα antagonist would be useful for the prevention and repair of inflammatory bone destruction and subsequent loss in the mouse model of CIA as well as human rheumatoid arthritis (RA) patients. This calls upon further clinical investigation to utilize its potential effect as an anti-arthritic drug.

## Introduction

Rheumatoid arthritis (RA) is a chronic autoimmune disorder that results in inflammatory joint damage aggravated by an array of inflammatory mediators like cytokines. TNFα is one of the primary cytokines which plays a critical role in the progression of various inflammatory diseases such as RA by regulating the production of IL6, IL1β, etc.[1]. Approaches to inhibit TNFα-induced inflammatory responses using monoclonal antibody inhibitors have therapeutic applications, but their use remained limited due to severe side effects [2]. Owing to the long halflife and thus prolonged inhibition of TNFα, monoclonal antibodies compromise the natural immune defense mechanism. These biologics, such as infliximab and etanercept are also reported to have less than 50% while adalimumab has less than 60% efficacy in RA patients [3]. The use of small-molecule antagonists is another approach for protein inhibition which prompted us to design a peptide that may lead to a more controlled intervention of the disease.

Peptides are active regulators and information agents making them suitable for drug discovery studies. Diversity in their chemical and biological nature along with high affinity, and specificity for molecular recognition are some of their virtues and advantages over conventional drugs. Lesser drug-drug interaction, minimal accumulation in the body, and much lower toxicity than other drugs are to name a few [4]. However, the limitation of peptides lies in the lower stability in the serum and if this issue is addressed then peptides may become more potent drugs.

This work focuses on designing and evaluating peptides in cell culture and collagen-induced arthritis (CIA) animal models for their anti-TNF and subsequent anti-arthritic activity. The peptide sequence PIYLGGVFQ (Mol. weight: 993.2 Da) was found to be most effective in computational screening followed by the cell culture assays. We found that this nonameric peptide showed significant inhibition of TNFα and was considered for its assessment in the CIA. It has a protective effect against joint damage as compared to untreated CIA mice.

## Materials and methods

### Selection of the peptide sequences followed by the docking studies

The TNFα structure was selected for peptide selection and docking studies as shown in Fig 1A, B. A single monomer subunit of the trimer was used for further studies. The potential binding site was extracted, using Acsite software (Fig 1C). This cavity was used in the docking studies of the peptides as described below. After docking was done with two different docking algorithms namely, Flex-X [5] and Genetic Optimisation for Ligand Docking (GOLD) [6], the interactive maps of the docked molecules were made in LIGPLOT [7] and the results were then analyzed to find the peptides that could bind with higher affinity to the TNFα and nine of the peptides were docked with varied binding energies. To confirm the results, docking was repeated with increased stringency by using a reduced cavity size as a receptor, which resulted in seven matching peptides which were again modeled with TNFα (Protein Data Bank, PDB ID: 1TNF) as a template using MODELLER software [8]. The scoring function, X-score was used to compare the binding energies [9]. The peptides were selected after visualizing them in Insight II. The peptides docked in a similar fashion, with binding energy within a cutoff of ±2.2 kcal/mol, and similar interacting residues of the receptor protein were shortlisted for solid-phase peptide synthesis. The selected hydrophobic peptide “PIYLGGVFQ” (after *in vitro* assays) as shown in Fig 1D was docked with the TNFα using Flex-X and GOLD (Fig 2A–D), and the interaction maps obtained from LIGPLOT (Fig 2E).

**Figure 1.**
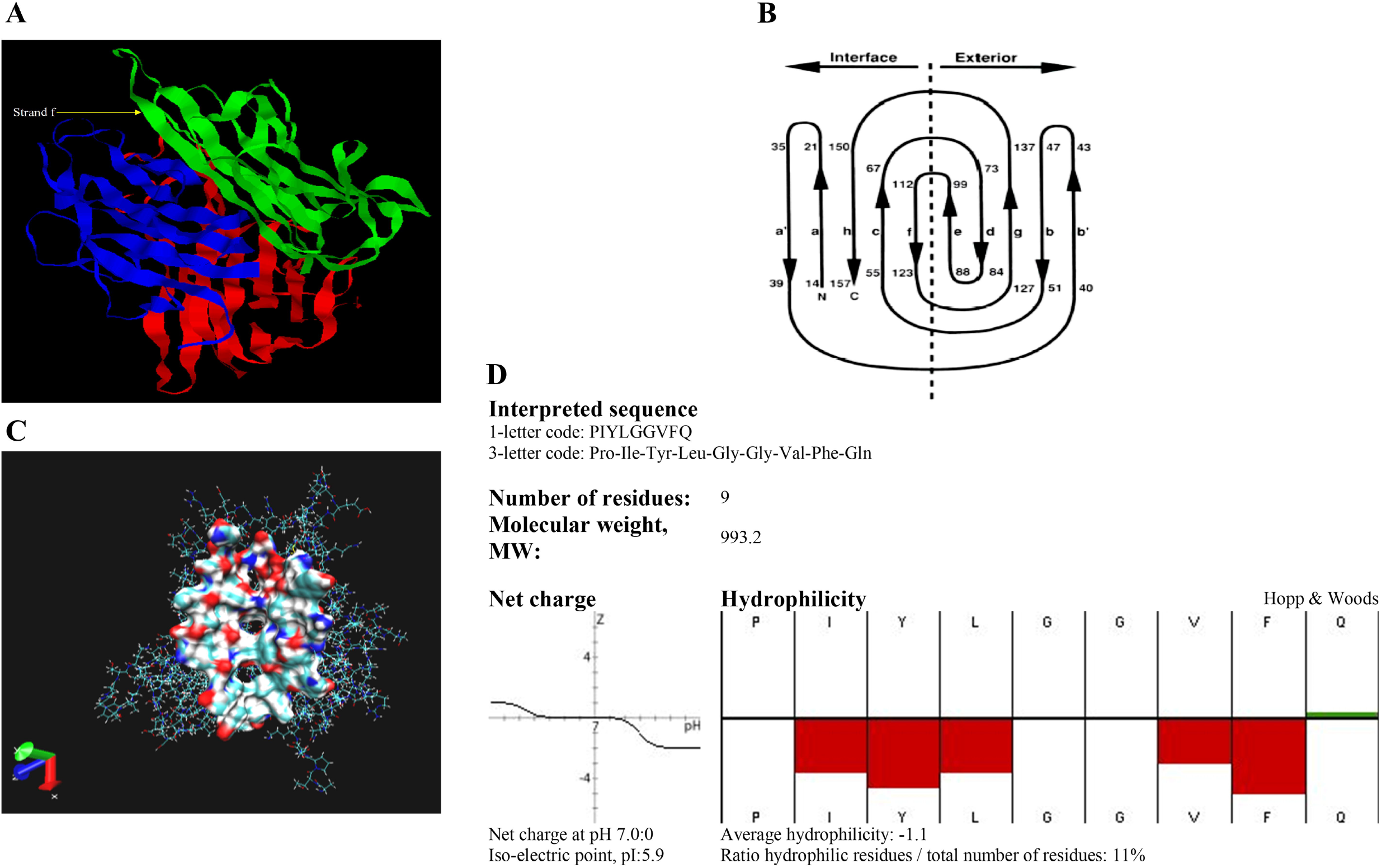
A. Structure of TNF (PDB: 1TNF). B. Topology diagram depicting the “jelly-roll” connectivity of the β-sandwich. C. Cavity extracted from the TNF trimer, using Acsite. D. Hydrophobicity plot.

**Figure 2.**
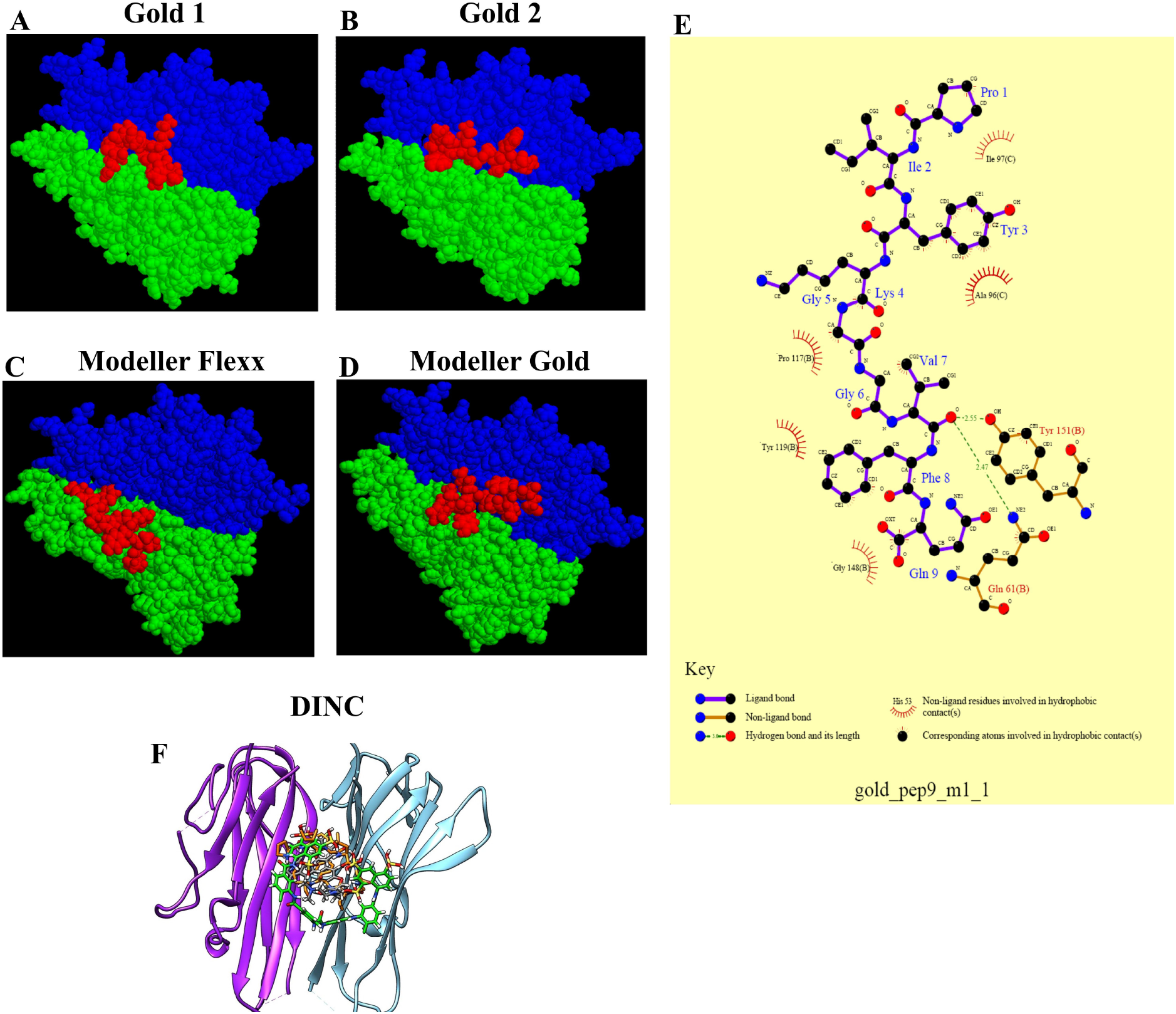
Docking studies show the TNFα monomers in blue and green structures and the peptide in red color, (A) Gold 1 (B) Gold 2 (C) Modeller Flexx, and (D) Modeller Gold. (E) Interaction map of the peptide with the TNFα, the interacting residues are also depicted here. (F) Docking pose of the co-crystalized ligand of PDB ID 2AZ5 (grey color sticks), Suramin (green color sticks), and peptide (orange color sticks) into the active site of TNF-alpha protein. Purple color ribbons represent chain A and blue color ribbons represent chain B of TNF-α.

The docking of peptides was also done through the Docking INCrementally (DINC) server [10] using peptide and TNF-alpha (PDB ID: 2AZ5) [11] as shown in Fig 2F. The peptide and protein structures were prepared in the Chimera program. For this molecular docking, the grid center X= −19.2, Y=74.5, Z= 33.8 Å, and box size 24X24X24 were selected around the co-crystalized ligand of the TNFα.

### Molecular dynamics (MD) simulation analysis

To validate the stability of the peptide inside the active site of TNFα, molecular dynamic simulation studies were done, using Assisted Model Building with Energy Refinement (AMBER)-12 software [12]. It provided information about the important binding interactions and free energy of the ligand, as well as the contribution of each residue to total binding interaction energies (Table 1). Protein structure (PDB ID: 2AZ5) has four chains: A, B, C, and D, where chain A-B is identical to chain C-D of the structure. Each A-B and C-D chains have the same bound co-crystallized ligand (307) in PDB (named as the standard, STD). As per some reports, A and B chains were used from PDB:2AZ5 for computational studies. Therefore A-B chain was used in to dock the peptide into the active site of TNFα [13]–[15]. Similarly, in MD simulation studies, chain A-B from the above structure was used as the peptide was docked into the active site of TNFα [11]. Since it is a structure obtained by X-ray diffraction studies of ligands bound to TNFα, STD was used as a reference molecule. This facilitated our understanding of the peptide binding mechanism (H-bonding, stability, and energy score).

Initially, the molecular docking studies were performed to place the STD and peptide molecules into the active site of TNFα, followed by the MD simulation. For the preparation of ligands and protein, the initial parameters and topology files (frcmod, prmtop, inpcrd, etc.) for the ligands (STD and peptide), protein, and protein-ligand complexes (TNFα-STD and TNFα-peptide) were generated using Antechamber [16] and tleap program of AMBER software by implementing, General Amber Force Field (GAFF) [17] and Amber ff99SB force field [18]. An explicit TIP3P water model was used to solvate the systems [19], and the solvation box was extended to 20 Å in all directions of the cubic box (solute forming). The systems generated were minimized followed by gradual heating from 0 to 300K, and density equilibration was done under the NPT ensemble. Then a constant pressure equilibration of 1 atm pressure (pressure relaxation time of 2.0 ps) for 1 ns at 300 K was applied. Subsequently, a 20 ns production run was done under NPT ensemble with a non-bonded interaction cut-off distance (12 Å). So, long-range electrostatic interactions were subjected to Particle-Mesh Ewald (PME) method [20], then the bulk effect simulation was performed by allowing periodic boundary conditions. The relative binding free energy was calculated for TNFα-ligand complex formation using the molecular mechanics-generalized born surface area (MM-GBSA) method [21] on the last four ns trajectory acquired from MD simulations of the protein-ligand complex formation to ensure conformational sampling and to obtain reliable binding free energy values. The Root Mean Square Deviation (RMSD) and B-factor analysis were performed with the CPPTRAJ program in AMBER [22], and the hydrogenbond occupancy analysis using the visual molecular dynamics (VMD) package [23].

After placing the known standard (STD) and our experimental peptide into the TNFα active site, the binary complexes (TNFα-STD and TNFα-peptide) formed after docking studies were subjected to MD simulation analysis for 20 ns. This was performed for the evaluation of the binding affinity of various ligands with the TNFα under multiple dynamic conditions. The study of the whole-system RMSD, ligand RMSD, protein-backbone RMSD, and B-factor/atomic fluctuations was done. The entire protein, and the protein backbone RMSD data, indicated that both binary systems were stable during the MD simulation study (Fig. 3A and B). The protein backbone and the whole protein were completely stable in binary complexes during the 20 ns simulation runs. The ligands were also found stable during the simulations especially after the 5 ns runs. The B-factor/atomic fluctuation data suggested that the fluctuations of atoms of the protein were very less during the simulations and most of the atoms/residues involved in atomic displacement were part of the loop region only (Fig. 3C and D).

**Figure 3.**
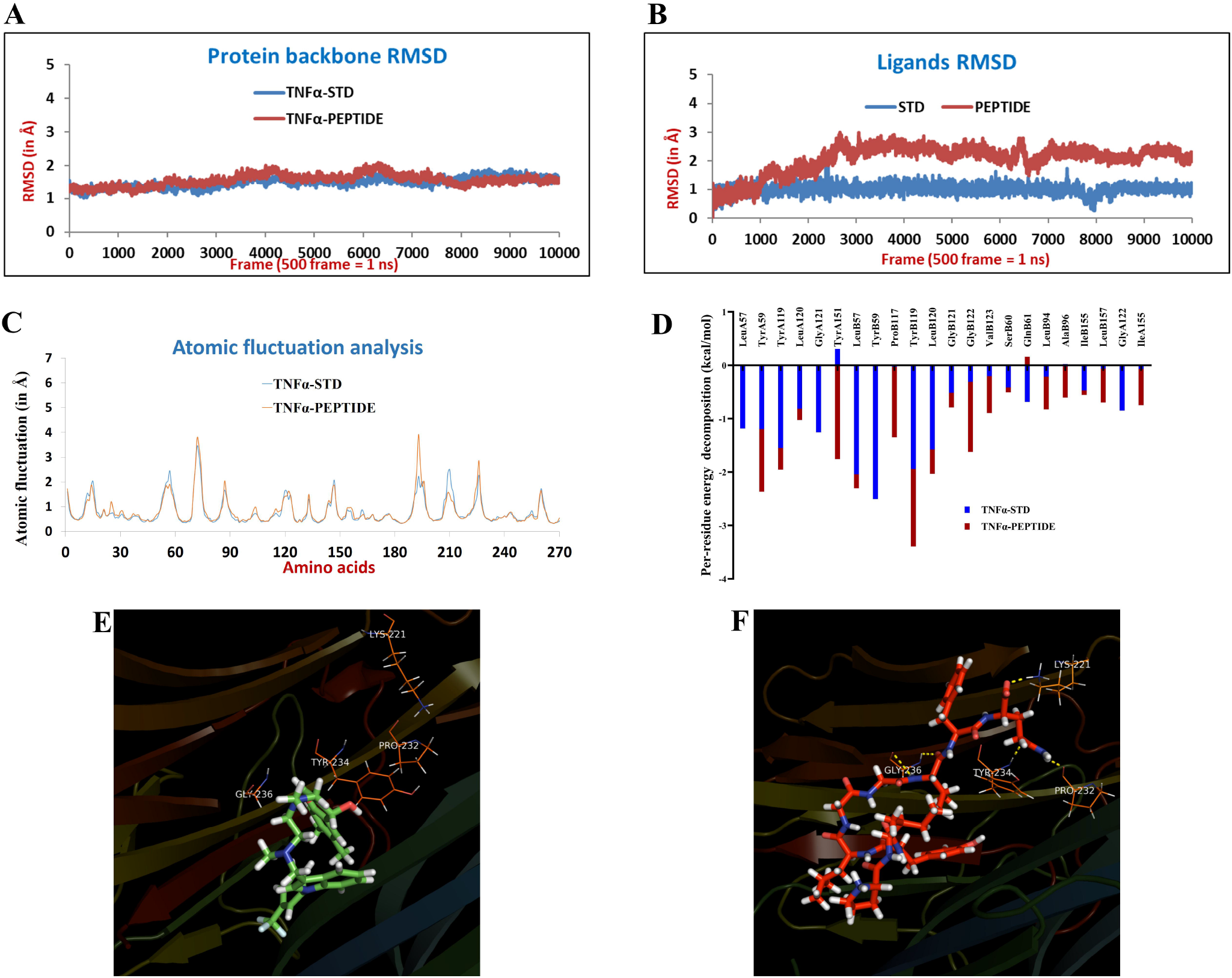
Molecular dynamics studies: (A) Protein backbone RMSD and (B) ligands RMSD in both binary complexes. (C) B-factor or atomic fluctuation studies and D) Per-residue decomposition analysis of the STD and the anti-TNFα peptide. The binding poses of (E) STD and (F) peptide docked into the active site of TNFα (PDB: 2AZ5).

The RMSD deviations for the receptor in both the binary complexes were ~0.75 Å and between the binary complexes was ~0.91 Å which are within the range of acceptability (0 ≤ 2 Å) and confirmed protein structure stability and shape in the dynamic state (Fig. 3A). The important active site residues (IleA58, LeuA57, TyrA59, SerA60, LysA98, ProB117, IleB118, TyrB119, GlyB121, GlyB122, TyrB151, etc.) were found to be identical in the binary complexes. The major RMSD deviation was observed for SerA86, ProB70, ArgB103, and minor RMSD deviation was noticed for GluA23, SerA71, CysA101, GluB23, SerB86, etc. The amino acid residues which were deviated in RMSD analysis were part of loop regions and found to be far from the core region of the active site. It was observed that no major RMSD deviations of ligands structure, either before or after the MD simulation studies were found in both binary complexes and were found to be less than 2 Å. This minor deviation shows that the ligands remained stable and consistently bound inside the active site of TNFα throughout the simulation runs.

It is well known that the active site of TNFα is completely hydrophobic and crystalized ligand (STD) also does not possess any hydrogen bond interactions with the key active site residues [11. A similar observation was noticed after the 20 ns MD simulation runs with STD. The designed peptide possesses various amino acids in its structure and due to the availability of various H-bond donors as well as acceptor centers, multiple H-bond interactions were observed during the molecular docking analysis (H-bond interactions with Lys98B, Pro117B, Tyr119B, and Gly121B) and similarly during the MD simulation. The H-bond occupancy results suggested that the H-bond interactions between protein and peptide were very weak. The per-residue decomposition analysis data suggested that Tyr119A, Leu57B, Tyr59B, Tyr119B, and Leu120B have a major contribution to the binding of STD in the active site of TNFα (Fig. 3D). While Tyr151A shows a slightly negative impact on STD binding. The residues Tyr59A, Tyr119A, Tyr151A, Leu57B, Tyr59B, Tyr119B, Leu120B, and Gly122B also show a greater contribution to the binding of peptides in the active site of TNFα. Figure 3E shows the binding poses of the STD with TNFα (PDB: 2AZ5) while the peptide docked into the active site has been depicted in Fig 3F.

The ΔG bind or the binding free energy contribution reported inhibitor (307/STD) demonstrated a strong binding affinity (−43.31 kcal/mol) while the designed peptide exhibits a comparable but lower binding affinity (−38.21 kcal/mol) for the TNFα protein.

### Synthesis and purification of peptide inhibitor for human TNFα protein

The synthesis was done by solid-phase peptide synthesis (SPPS) process using the Fmoc strategy. The nature of the peptides essential for their purification using the peptide calculator (http://www.innovagen.se/custom-peptide-synthesis/peptide-property-calculator/peptide-property-calculator.asp) showed that the peptide is highly hydrophobic (Fig 1D). After SPPS, they were cleaved and desalted on the LH-20 column. Then the peptides were dissolved in methanol and concentrated with rotavapor followed by lyophilization. The yield of the peptide was: between 60 and 80 mg. The peptides were purified using reversed-phase HPLC (Fig S2C). Briefly, 25 μl of the peptide with appropriate dilution was injected and resolved with the mobile phase (methanol). The identity and molecular weights of the peptides were confirmed by mass spectrometry. The micromolar quantity of the peptide was dissolved in 0.1% TFA. The solution was mixed with an equal amount of matrix 4-hydroxy cyanocinnamic acid (HCCA), prepared in acetonitrile and 0.1% TFA (1:2). The matrix-analyte sample (1 μl) was loaded on the steel plate and left for crystallization for 5 min. The compound was then analyzed by MALDI-TOF/TOF mass spectrometer. The mass of all the peptides matched the calculated mass and the peak was clean and single (Fig.S2B).

### Cytotoxicity assay

The Wehi-164 murine fibrosarcoma cell line undergoes apoptosis by TNFα stimulation in the presence of actinomycin D (AcD), which makes it suitable for such an experiment [25]. On the other hand, immune cells such as J774 macrophages, Jurkat cells, etc., show cell proliferation in response to TNF-α. So, in this study, 70-80% confluent Wehi-164 cells, with more than 90% viability were used. In 96-well plates, 5×10^4^ cells per well were taken in the presence or absence of AcD, 0.2μg/ml, TNFα (100 ng/ml), peptide, and suramin (100 and 200 μM), premixed at 4°C for 1h. After 20h incubation at 37°C, 25 μl of MTT solution (5 mg/ml in 1X PBS) was added and incubated for 4 h. Then isopropanol (with 0.04 M HCl) was used to dissolve the formazan before measuring the OD at 570 nm with background subtraction at 650 nm (Molecular Devices, USA) to determine the TNFα-induced cytotoxicity.

### Preparation of nuclear extract

The cell nuclear protein was extracted using the method of Schreiber and coworkers [26] with a minor modification. Briefly, A549 cells at a concentration of 10^6^ per well were incubated with or without TNFα (100 ng/ml) and different concentrations of peptide or suramin, for 45 min followed by washing with PBS. These treated cells were then resuspended in the cell lysis buffer [10 mM KCl, 10 mM HEPES (pH 7.5), 0.1 mM EDTA, 0.5% NP40, 1 mM DTT, and 0.5 mM PMSF with mammalian protease inhibitor or PI (Sigma)]. The cells were kept in ice for 25-30 minutes with intermittent mixing, then vortexed followed by centrifugation at 12,000 g for 10 min at 4°C. The pellet containing the nuclei was washed with the cell lysis buffer before resuspending in the nuclear extraction buffer (0.4 M NaCl, 20 mM HEPES with pH 7.5, 1 mM EDTA, 1 mM PMSF, 1 mM DTT with PI solution) and incubated in ice for half an hour. The nuclear protein extract was then collected as the supernatant after centrifugation (12,000 g at 4°C for 15 min).

### Electrophoretic Mobility Shift Assay (EMSA)

A dsDNA probe for NFκB (Promega) was used for the gel shift assay after labeling with radioactive ATP [γ-32P] by T4 polynucleotide kinase using the manufacturer’s instructions. The reaction mixture contained 2 μ1 of 5x binding buffer (50 mM Tris–HCl (pH 7.5), 20% glycerol, 250 mM NaCl, 5 mM MgCl_2_, 2.5 mM DTT, 2.5 mM EDTA, and 0.25 mg/ml poly-dI-dC) and 2 μg of nuclear extract and nuclease-free water. A probe (1 μ1, 10^6^cpm) was added to initiate the reaction for 30 min at room temperature. The protein binding specificity with the DNA was assessed by competition reactions, where 20-fold molar excess of oligonucleotides (unlabeled) was added to each tube prior to adding a radio-labeled probe. All the prepared samples were subjected to native gel electrophoresis (4% polyacrylamide gel). The radiographic gels were scanned (Fuji FLA-2000 Phosphor imager) for further analysis.

### Western blot analysis

The nuclear protein extracts from the TNF-stimulated A549 cells were subjected to SDS-PAGE and transferred to a nitrocellulose membrane (Millipore, USA) using the transfer buffer (192 mM glycine, 25 mM Tris, 20% methanol) for 20 min at 20 V semi-dry transfer apparatus (BioRad, USA). Blocking was done with 1% BSA for 1 hour at room temperature. After three times (5 min each) washing with 1X TBST, the membrane was incubated overnight with anti-NFκB (p65) and anti-β actin antibodies at 4°C. It was then probed with anti-rabbit IgG HRP for 1 h after the TBST wash. The blot was washed again and developed using 4-chloronaphthol (Sigma). The band intensities were measured and the densitometric analysis was performed with the Alpha Digi Doc software tool (Alpha-Innotech Corporation, USA).

### Immunocytochemical assay

Inhibition of the TNFα-induced NFκB activation resulting in NFκB RelA/p65 nuclear translocation in A549 cells by the peptide was visualized by immunocytochemistry assays. The A549 is a human lung epithelial cell line that responds well to the stimulation by TNF-α resulting in the NFκB activation [25]. The cells were seeded at a concentration of 2.5 x 10^4^ cells/well and cultured overnight before incubating for 45 min with TNFα (100 ng/ml) with and without the peptide (premixed for 1 hour). The cells were then washed with isotonic buffer PBS followed by fixing and blocking (PBS with 0.05% sodium azide and 1% BSA) for 30 min and the cell permeabilization was done with Triton x-100 (1%) for 30 min. For the nuclear staining, 10 μl of propidium iodide (0.5 mg/ml) was mixed in the well and incubated at 4°C for 30 min in dark.

The cell staining for p65 was done using the supplier’s protocol (Santa Cruz, USA). Briefly, cells were incubated in the primary antibody against NFκB p65 (Santa Cruz, USA, sc-372) diluted in blocking buffer (overnight, 4°C), followed by FITC-conjugated secondary anti-rabbit IgG (1:200) was added to the cell and incubated for 1 hour. Cells were washed thrice with PBS and scanned under a fluorescence microscope (Nikon, Japan).

### Inhibition of cell surface binding of TNF-α

The U937 cells were plated at a density of 2×10^4^ cells per well (96-well plate) and stimulated with human TNFα (100 ng/ml) with and without the anti-TNFα peptide for 1 hour at room temperature. Then the cells were treated with the primary and FITC-labeled secondary antibodies as described above. The TNF-α binding to receptors was analyzed by fluorescence microscope and image processing was done with ImageJ (NIH).

### Analysis of LPS-induced cell surface TNFα by flow cytometry

The peptide decreases the TNFα signaling through NFκB and therefore it should decrease surface TNFα as well. As an indirect approach to evaluate the effect of the peptide to quantitate surface TNFα with and without the peptide, the expression of surface TNFα was measured by flow cytometry. For this experiment, U937, a pro-monocytic, human myeloid leukemia cell line, was used because it expresses the receptors of TNF-α (TNFR1 and TNFR2) in a substantial number [24]. For all flow cytometry experiments, LPS (1 μg/ml) was dissolved in RPMI. The peptide was dissolved in DMSO (1% v/v) before diluting with RPMI. The U937 cells were transferred into microcentrifuge tubes at a concentration of 4×10^6^/ml. The cells were treated with 1 μg/ml LPS with or without the peptide (200 μM) for 2 h. Cells were then resuspended in 200 μl FACS buffer. All the cells were labeled with monoclonal anti-human TNFα antibody (Peprotech, USA) and subsequently washed with 1X PBS followed by treatment with FITC-conjugated secondary anti-mouse antibody. After the wash flow cytometry was performed. A total of 50,000 events were collected using a FACS Calibur flow cytometry system (BD Biosciences, USA), and data were analyzed using the FlowJo program.

### Animal experiments

Animal experiments were performed according to the institutional guidelines and the protocol was approved by the animal ethics committee of the National Institute of Immunology (NII), New Delhi. A total of 20 female DBA1/J, 6 to 8 weeks old mice were procured from the inbred facility of NII. Mice were housed in up to 6 mice per cage in a room maintained at 23 ± 2°C with 50 ± 10% humidity and 12-h light/12-h dark cycles. The animals were allowed free access to tap water and regular rodent chow.

### Induction of CIA and treatment

Induction of CIA was done in accordance with our previous study [27], with some minor modifications. Briefly, the mice were acclimatized for 7 days and divided into four groups, healthy, CIA, CIA+peptide (5 mg/kg), and CIA+peptide (10 mg/kg). Thereafter, all mice except the healthy group were immunized by intradermal injections of an emulsion containing 100 μg of immunization grade bovine type II collagen (CII) in the Freund’s complete adjuvant at the base of the tail. On day 21, a booster containing 100 μg CII emulsified with Freund’s incomplete adjuvant was administered and observed for disease development. Treatment was initiated after 12 days of the booster dose when the disease severity of all groups was observed to be the maximum. The peptide was dissolved in DMSO (≤ 1% v/v) followed by dilution in 1X sterile PBS before administering intraperitoneally (i.p) in the mice, three times a week. The healthy group was treated with a vehicle (1% DMSO in PBS (v/v)) alone. A schematic presentation of the disease induction and treatment protocol is shown in Fig 5A. All the groups of mice were regularly scored for disease indicators (inflammation, redness, severity, etc.) according to their severity until termination. All the reagents used for the generation of the CIA models were obtained from Chondrex (USA).

**Figure 4.**
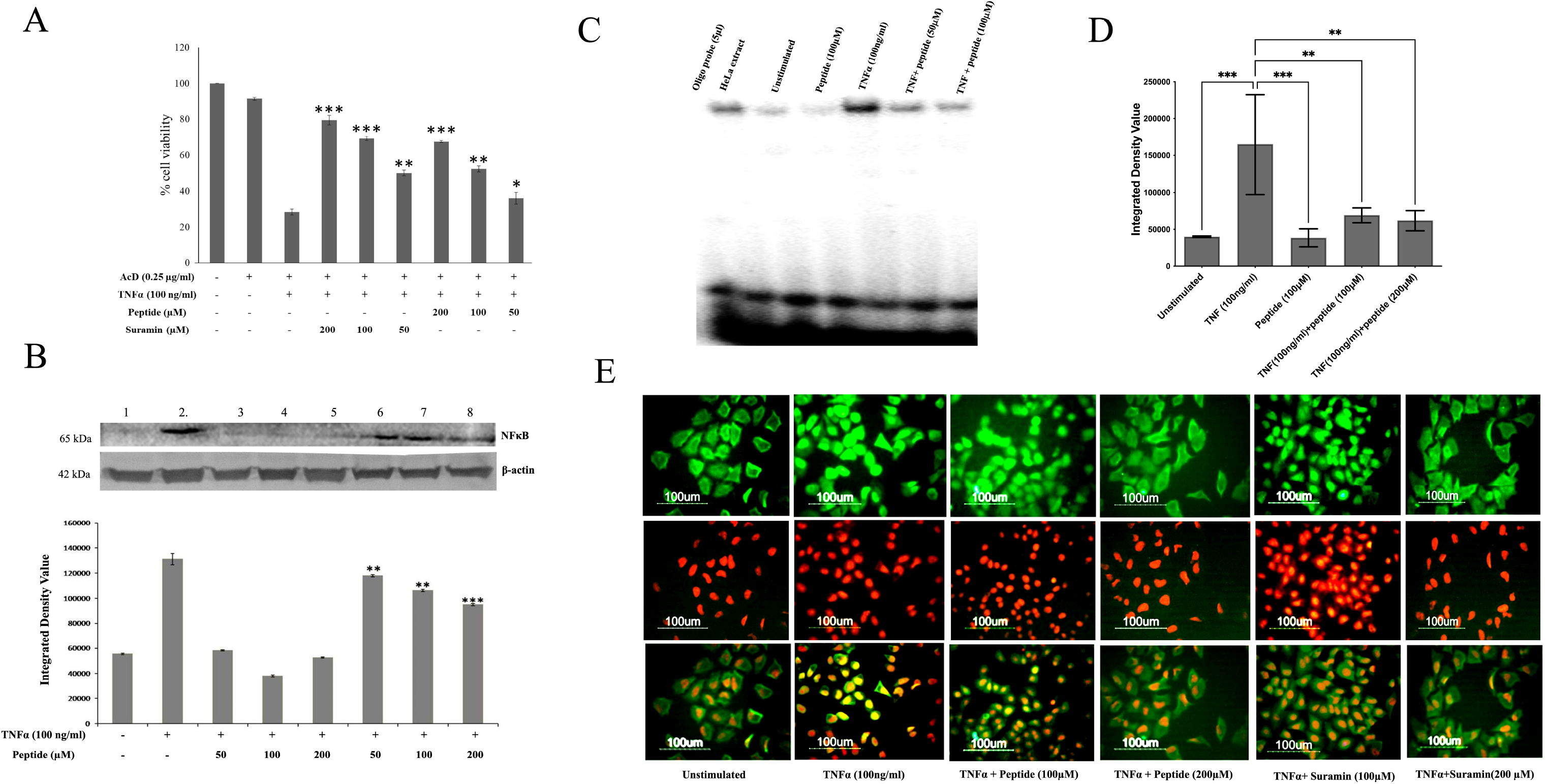
Cell culture assays for *in vitro* evaluation. (A) Cytotoxic effects of the TNFα on Wehi-164 cells and the protective effect of the peptide and suramin as evaluated through MTT assay. (B) Western blot of nuclear extract after TNFα induced NFκB translocation in A549 cells and its inhibition by the peptide at the concentrations of 50, 100, and 200 μM. The constitutively expressed protein β-actin was found to be expressed equally. (C) EMSA of nuclear extracts (20 μg of nuclear extract of the unstimulated, TNFα stimulated cells with or without the presence of peptide. (D) The densitometric analysis (IDV) of the EMSA bands (n=3) (E). The immunocytochemical assay shows the inhibitory effect of peptides (100 and 200 μM) on TNF-mediated nuclear translocation of TNFκB in the A549 cell line.

**Figure 5.**
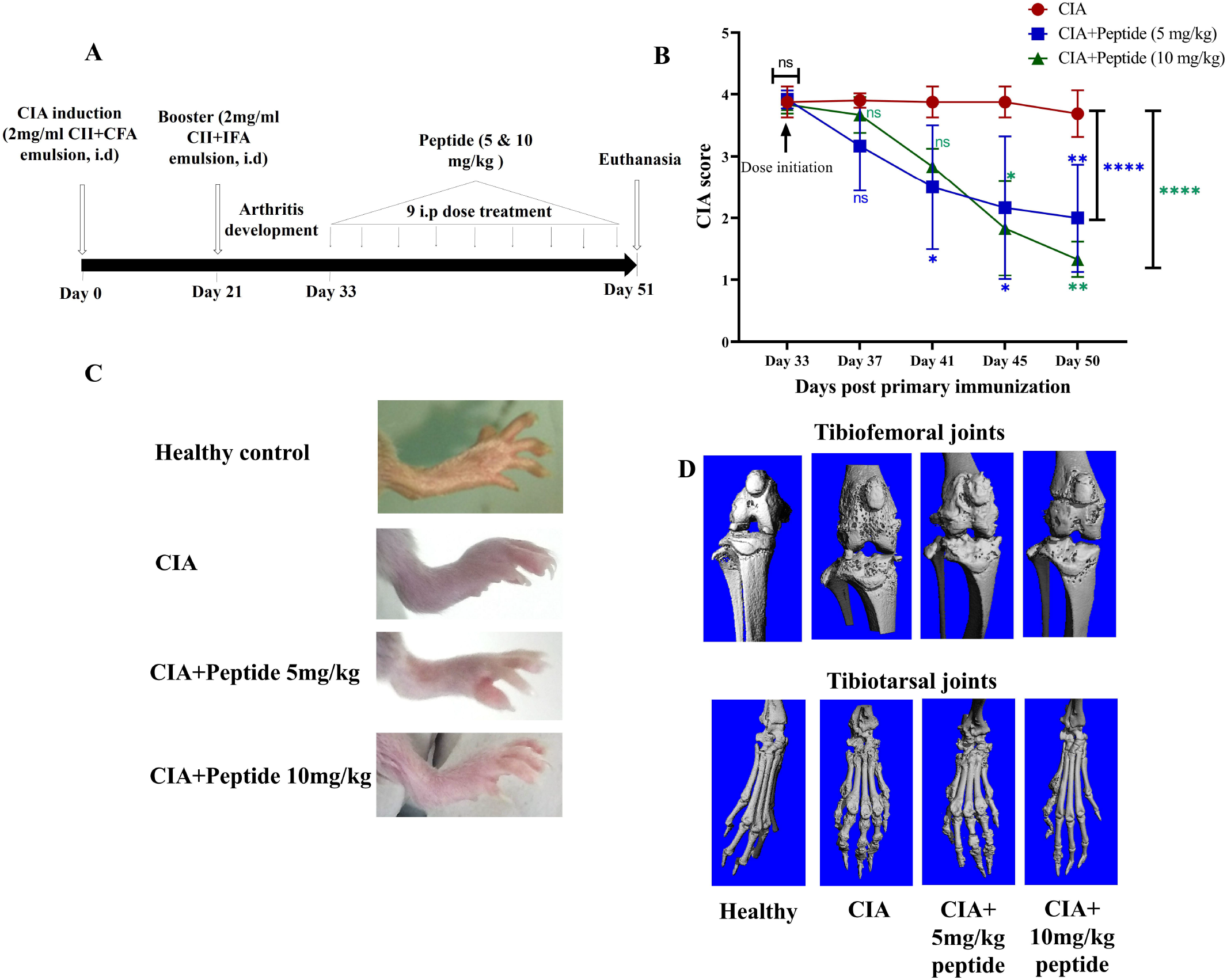
*In vivo* evaluation studies in CIA animals. (A) The elaboration of CIA induction protocol, treatment regimen, and termination of the experiment. (B) The plot shows the disease progression after the initiation of peptide treatment at 5 mg/kg (blue) and 10 mg/kg (green) in comparison with the untreated group (red). (C) The morphological appearance of the mice’s hind paws. (D) Micro-CT images show the comparative account between the experimental groups. * p≤ 0.05, **p≤ 0.01, ***p≤ 0.001, ****p≤ 0.0001.

### Clinical evaluation of the disease

To assess the disease characteristics, all the mice were scored for disease extent at 3-4 day intervals for ~20 days following the booster immunization as described in our previous study [28]. Both paws and joints of fore and hind limbs were scored as follows: 0: Normal; 1: redness and swelling of the paw or one digit; 2: swelling and redness of ankle and wrist; 3: severe redness and swelling of entire paw including digits and 4: severe arthritis of the limb including entire paw and digits and expressed as the mean of all 4 limbs of a mouse.

### Micro-CT scanning and image processing

Micro=CT scanning of hind-limbs from healthy control, CIA, CIA+peptide (5mg/kg), and CIA+peptide (10mg/kg) groups were performed at 70kVp, 114μA by exposing the bones to radiation for 300ms. The medium used for scanning was formalin. Scanning at 10μm resolution was performed using a 0.1 aluminum filter to reduce the beam hardening effect. Once the scan was completed, the in-built reconstruction algorithm was used to develop the cross-sectional images. The range of global threshold, 654.2-1239.7 mg HA/ccm, was used to develop the 3D geometry of the bones.

### Statistical analyses

The statistical data analyses were performed using GraphPad Prism software. Three experimental values(n) are given as mean ± standard deviation (S.D.) in the cell culture assays. The student’s *t*-test was employed to calculate the differences from the respective controls for each paired experiment. The data from animal experiments have been expressed as mean ± SD (n = 5 or 6). As applicable, *p*-values were calculated both using unpaired t-tests, and one-way or two-way ANOVA tests; and *p* ≤ 0.05 was significant.

## Results

### *In silico* designing of a peptide inhibitor of human TNFα

The structure of TNFα in solution is a homotrimer containing monomers of 17,350 Da each as determined by Eck et al (Fig 1A). The monomers have a ‘jelly-roll’β-sheet (anti-parallel) sandwich structure composed of ten β-strands [29]. As shown in fig 1B, all the a, h, c, and f strands form a flat inner sheet of the ‘jelly roll’ which are involved in monomeric strand contacts during the trimer formation, whereas, the carboxy and amino strands are stacked together in the inner sheet. The outer sheet which is a highly curved structure is composed of the strands b, g, d, and e thereby forming the outer surface of the TNFα trimer. The strand b present in the outer sheet is interrupted, an unusual phenomenon in β-sandwich structures; at the amino terminus, which is residue 26, a chain of twenty (20) amino acids forms the strands a1 and b1. These strands provide stability to this structure by surrounding the 2 sheets at the NH_2_-terminal edge. In a TNFα monomer, there are 3 helical segments, while none of these helices extend more than one full term. The amino acid stretches involved in the formation of these helices include 106 to 110, 138 to 142, and 145 to 150. Also, each monomer contains a disulfide (-S-S-) bridge serving as a connection between one loop (connecting strands c and d) with the other loop (connecting strands e and f). The amino acids at 69 and 107 positions together form this disulfide bond [29].

### Identification of peptide sequence

Fourteen peptide stretches were selected (Table S1) and synthesized *in silico* for docking studies. The interacting stretches in each monomer of TNF were located, using the Database of Interacting Proteins (DIP), and the results were confirmed with Protein-Protein Interaction (PPI) server (Tables S 2, 3, 4, 5, 6, and 7). All the results were compiled and the stretches coming at the interface were marked as can be viewed in Figure S1.

### Docking studies of peptides

The molecular docking studies were done in the DINC server using the co-crystallized ligand of PDB:2AZ5 to check whether the DINC docking program can generate the same binding pose as found in X-ray data (Fig 2F). It was observed that the co-crystallized ligand of TNFα also had the same binding pose. Similarly, suramin and the peptide were also docked into the active site. In these studies, suramin (−10.60 kcal/mol) and co-crystalized ligand (−10.10 kcal/mol) exhibited almost the same binding energy. The peptide exhibited the −8.00 kcal/mol binding energy among the other ligands. This suggests that although the peptide is loosely bound to the TNFα as compared to the other molecules, it has a comparative binding affinity with the gold standard control molecules.

### Peptides inhibiting TNFα induced apoptosis in Wehi-164 cell lines

Wehi-164, a TNFα-sensitive cell line was incubated with TNFα for 20 hours in the presence of actinomycin D (AcD) which is an NFκB transcription inhibitor resulting in apoptosis (Fig. 4A). Three different concentrations of the peptide (50, 100, and 200μM) in the presence of 100 ng/ml TNFα and AcD (0.25 μg/ml) suppressed apoptosis significantly and in a dose-dependent manner (*P≤0.05, **P≤0.01, ***P≤0.0001). The costimulated cells with TNFα and AcD showed maximum cell death (70%) with respect to the unstimulated cells. The peptide at 200 μM concentration showed a significant reduction in apoptosis (26.4%). Suramin, a known anti-TNF compound showed inhibition with 50.44% and 43.5% apoptosis at 200 and 100 μM concentrations. Cells incubated with the peptide at 200 μM concentrations showed low cytotoxicity.

### Inhibition of NFκB activation and nuclear translocation in A549 cells

The level of the NFκB (p65) in the A549 nuclear extract was measured in the TNFα stimulated cells, and the effect of the peptide was also assessed using Western blotting (Fig 4B). The protein level of the housekeeping gene β-actin was used as a positive control for the nuclear extract. It was shown that the peptide at the increasing concentrations (25, 50, and 100 μM) significantly inhibited (45%, 46%, and 55% respectively) the NFκB nuclear translocation.

The inhibition of the nuclear translocation by the peptide was also confirmed through the gelshift assay or EMSA (Fig 4C). The quantitative analysis of each band of EMSA is represented as integrative density values (IDV) in Fig 4D. The addition of an excessive unlabeled oligonucleotide probe prevented the band shifts while exhibiting the protein-DNA interaction specificity.

### Immunocytochemical analysis of the nuclear translocation of NFκB in A549 cells and TNFα localization on the cell surface of U937 cells

The visualization of the NFκB nuclear translocation in the A549 cells was performed using the immunocytochemical localization assay (Figure 4E). In TNFα stimulated cells, the nuclei were observed to be fluorescent green, unlike the unstimulated cells where the nuclei looked hollow. An orange stain propidium iodide (PI) was used to locate the nuclei of the cells. The peptide showed almost similar inhibition of NFκB activation hence the nuclear translocation. As a control, we have used a TNF-α-specific inhibitor, suramin in our experiments. Suramin was used to compare the anti-TNF activity of the peptides using TNF-induced cytotoxicity in Wehi-164 cells (Fig 4A) and the inhibition of TNF-induced NFκB nuclear translocation in A549 cells as shown in the immunofluorescence assay (Fig 4E). Suramin was used along with the peptides showing similar inhibition of NFκB nuclear translocation in the immunofluorescent cells. These results were consistent with the EMSA as well as MTT assay results.

The human monocyte cells, U937 were incubated with TNFα with or without the peptide and were visualized under the fluorescence microscope to observe the localization of the green color depicting the presence of TNFα. Fluorescent green periphery (Fig S3) was observed in TNFα stimulated cells however in the unstimulated cells the color was diminished and not localized, while the peptide incubated cells also showed lower FITC-antibody localization on the periphery, the intensity of which lies between the unstimulated and the TNF-only stimulated cells. This suggests the inhibitory activity of the peptide on the TNFα.

### The use of peptide inhibitor decreases cell surface expression of TNFα in LPS-treated cells

The LPS induced TNFα production, and its subsequent binding on the U937 cell surface was inhibited by the peptide. The decreased nuclear translocation of activated NFκB in A549 cells also assessed the inhibition.

The LPS-stimulated U937 cells showed a higher presence of TNFα on the cell surface as shown by the proportion of the histogram peak towards the right of the gate (blue line) (Fig S4B). The amount of non-fluorescence cells represents a lower presence of TNFα on the cell surface and is shown towards the left of the gating blue line. With the LPS treatment, a higher proportion of the cells are shifted right (Fig S4B) while the presence of the peptide in the LPS-stimulated cells showed a shift more towards the left, which is an intermediated stage between the unstimulated (Fig S4A) and the LPS-stimulated cells alone (Fig S4C). In the LPS-treated group, TNFα surface expression was markedly above the unstimulated cells. Treatment with 200 μM peptide significantly inhibited LPS-induced TNFα surface expression versus LPS only in both experiments. This suggests that the peptide causes the inhibition of TNFα trimer formation, this trimer is essential for its binding to the TNF receptors. The decreased binding is thus detected in the peptide-treated cells.

### Peptide treatment suppresses CIA score in mice

The arthritic symptoms appeared after the injection of booster immunization on day 21, depicted in the experimental outline (Fig 5A). After 12 days of post-booster dose, the symptoms were found to be severe. In the CII-immunized mice periarticular erythema, edema, and functional loss in gait were observed. The walking pattern of the arthritic animals improved markedly in all the peptide-treated groups as opposed to the untreated CIA. The peptide treatment (5 and 10 mg/kg) showed a substantial decline in the mean arthritic score compared to the healthy control. There was a dose-dependent reduction from day 16 post booster, with the CIA score pattern similar in both the 5 and 10 mg/kg peptide doses. A decline in CIA score in 10 mg/kg from day 41 (p ≥ 0.05) to day 50 (p ≤ 0.01) was observed to be significant throughout (Fig 5B). The overall CIA score difference was also calculated as a cumulative effect, on the experiment termination day (day 51), which showed highly significant improvement (p< 0.0001****) in both the peptide 10 mg/kg group and the 5 mg/kg group. The therapeutic effect of the peptide can also be observed in pictures of the hind limb footpads (Fig 5C). These results corroborated with that of the CIA scores.

### Peptide treatment inhibits CIA-induced joint damage

MicroCT analyses exhibited protective effects of the peptide. The hind limb microCT pictures of the untreated CIA animals showed joint erosion with osteophyte formation (Fig 5D). While, with the treatment of peptides 5 mg/kg and 10 mg/kg, a marked recuperation of the joint damage was observed in both knee (tibiofemoral) and ankle (tibiotarsal) joints of the mice. The knee joints were substantially improved in both the treated groups. However, both the peptide-treatment groups showed joint damage as compared to the non-immunized mice.

## Discussion

Rheumatoid arthritis (RA) is a T-cell-mediated disease, however, pro-inflammatory cytokines are prevalent in the synovium and plasma of RA patients [30]-[32]. One of the prevalent therapeutic approaches in RA is to inhibit the proinflammatory cytokines. TNFα is known to be a prime cytokine because it stimulates the production of IL-6, IL-1, and GM-CSFs in the synovium, it increases the level of an osteoclast differentiation factor, the RANKL thus aggravating the lesion formation in RA [33]. It is now widely reported that the TNF-neutralization with monoclonal antibodies or soluble TNF-receptors markedly prevents or even repairs the inflammatory bone damage in RA [33]. Many TNFα inhibitors such as infliximab, etanercept, and adalimumab have been approved by the US FDA and are used in clinical practice [34]. However, there are concerns with the use of injectable protein-based TNFα inhibitors that may also be addressed with small-molecule inhibitors. Overuse of the former has been associated with an increased incidence of latent tuberculosis. The drugs, etanercept, and infliximab have been reported to show a risk for congestive heart failure. In addition, a potentially increased incidence of lymphoma has been observed in patients treated with the injectable TNFα blocking agent, including adalimumab [35]. In our study, we showed that the inhibitory effects of a TNF-deactivating peptide not only antagonize inflammation but also protect against joint destruction in the autoimmune disease model (Figure 5D). The molecular weight of one of the anti-TNF antibodies adalimumab is almost 149 times more than the peptide under consideration, which makes the latter a potentially effective drug candidate. As a result, there would be minimal drug-drug interaction, less accumulation in the tissues, and lower toxicity. The downside of using peptides is attributed to their instability in the physiological environment and limited membrane permeability. Other features to be addressed are the proteolytic degradation in blood limiting the peptide’s bioavailability and half-life. To overcome these, an increase in stability can be incorporated using various biomaterials, and frequent dosing to maintain the physiologically effective serum concentration [36].

There are many small molecules under different stages of preclinical and clinical trials that inhibit protein synthesis, such as thalidomide which is a p38 MAP kinase inhibitor, and TACE inhibitors. Many of them have quite successful and promising anti-inflammatory activity by inhibiting IFNγ, IL-12, etc. Such property may be hazardous as it may cause immunosuppression of the recipient [37]. Like RDP1258 along with its D-isomer, Allotrap 1258, has enhanced protection from cytotoxicity of T lymphocytes [38], functional peptides suppress the IκB degradation causing blockade of NFκB transcription [39]. A peptide WP9QY which simulates the contact site of TNF-receptor and ligand rescues from bone resorption by obstructing the osteoclast recruitment and activation by RANKL [40]. However, such peptides have been reported to show a high and non-specific inhibition of their respective targets causing adverse events.

In the present study, we developed a small peptide inhibitor of TNFα. The stimulated macrophages produce membrane-bound 27kd TNFα that first trimerizes, and then can either bind directly to TNFR-55 and TNFR-75 receptors through cell-to-cell contact or undergo cleavage by TNFα converting enzyme (TACE) and bind in its soluble form. Thus, trimerization is one of the most important steps for TNFα activity and was targeted for designing peptide inhibitors. This may be attributed to the TNF-TNFRII interaction. Because, studies have found that TNFRII signaling results in a protective effect in inflammatory arthritis, while TNF-TNFRI results in proinflammatory activity [41]. The TNF ligand-receptor (TNFRII) blockage has been reported to worsen the arthritis symptoms[42].

With the TNFα administration, in absence of actinomycin D, the cell viability was observed to be increased. Although TNFα is an apoptosis-inducing factor, the TNF receptor contains the death domain-like Fas [43]-[45]. However, in most cases, it does not induce apoptosis because TNFα induces activation of NFκB, which inhibits apoptosis. The loss of NFκB inducibility pronounces a decrease in the cell viability [46], [47]. Our peptide prevented the TNF-induced activation of NFκB thus effectively inhibiting the nuclear translocation of p65. This anti-TNF peptide was further evaluated using animal studies for its potential use as a peptide drug. The decrease in viability due to TNFα was significantly inhibited by our peptide. After performing the cell culture assays, we tried to assess its efficacy in the inflammatory animal model. The peptide sequence was subjected to Protein BLAST (https://blast.ncbi.nlm.nih.gov/Blast.cgi?PROGRAM=blastp&PAGE_TYPE=BlastSearch&LINK_LOC=blasthome) to see the sequence similarity with *Rattus norvegicus* and *Mus musculus*, interestingly, we could not find any similarity with rat however there was complete sequence similarity with the mouse species. This finding encouraged us to assess the antiarthritic activity of the peptide using the CIA mouse model. The peptide showed significant protective activity on the autoimmune disease in the animal. TNFα is known to induce osteoclastogenesis, therefore, its inhibition prevents bone resorption thus protecting the periarticular bone in RA [48], which is in concurrence with our micro-CT results. Even though histopathological studies were not performed for the *in vivo* study, the micro-CT analysis results strengthened our belief in a peptide-mediated reduction in the arthritic activity of TNFα. We acknowledge that the use of a scrambled peptide would have strengthened our study. However, it can be hypothesized that the peptide inhibits bone resorption by indirectly working against RANKL in the CIA mice. To conclude, the PIYLGGVFQ peptide was successful in inhibiting the inflammation at *in vitro* cellular level and *in vivo* by inhibiting bone destruction. This peptide will be a valuable prototype for the development of small molecules for the prevention of inflammation and bone or joint damage deactivating TNFα and the associated inflammatory mediators.

## Acknowledgment

The authors acknowledge all the lab members for insightful discussions and critical proofreading. We also acknowledge the animal research facility staff of the National Institute of Immunology (NII), for the care of animals and support for conducting experiments.

## Author Contributions

DS, CG, HRD, and CJL have conceptualized and designed the experiments; DS, CG, and SS performed the cell culture experiments; RY, PG, and VKS performed the *in silico* study design and analyses; S Kashyap, S Kumar, VPD and AKP facilitated and performed the animal experiments. DS, CG, SH, PG, VKS, and CJL contributed to the writing and reviewing of the manuscript.

## Ethics approval

All animal studies were performed as per the institutional guidelines and the protocol was approved (IAEC#367/15) by the Institutional Animal Ethics Committee (IAEC) of the National Institute of Immunology (NII), New Delhi, India.

## Figure legends

**Figure S1.**
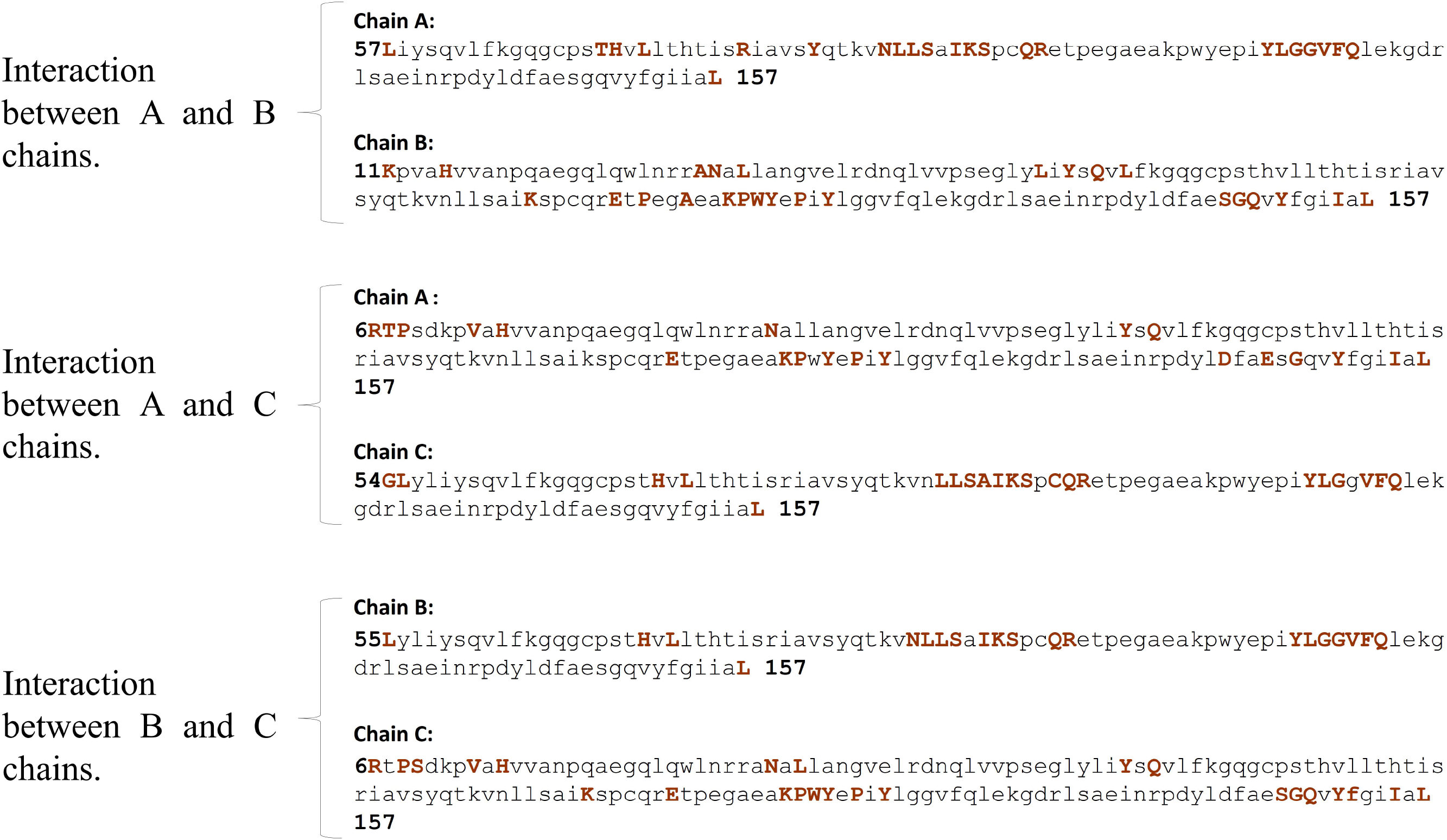
The stretches of residues are present at the interface of TNF monomeric chains. The residues involved in the interaction are shown in colored and upper-case letters.

**Figure S2.**
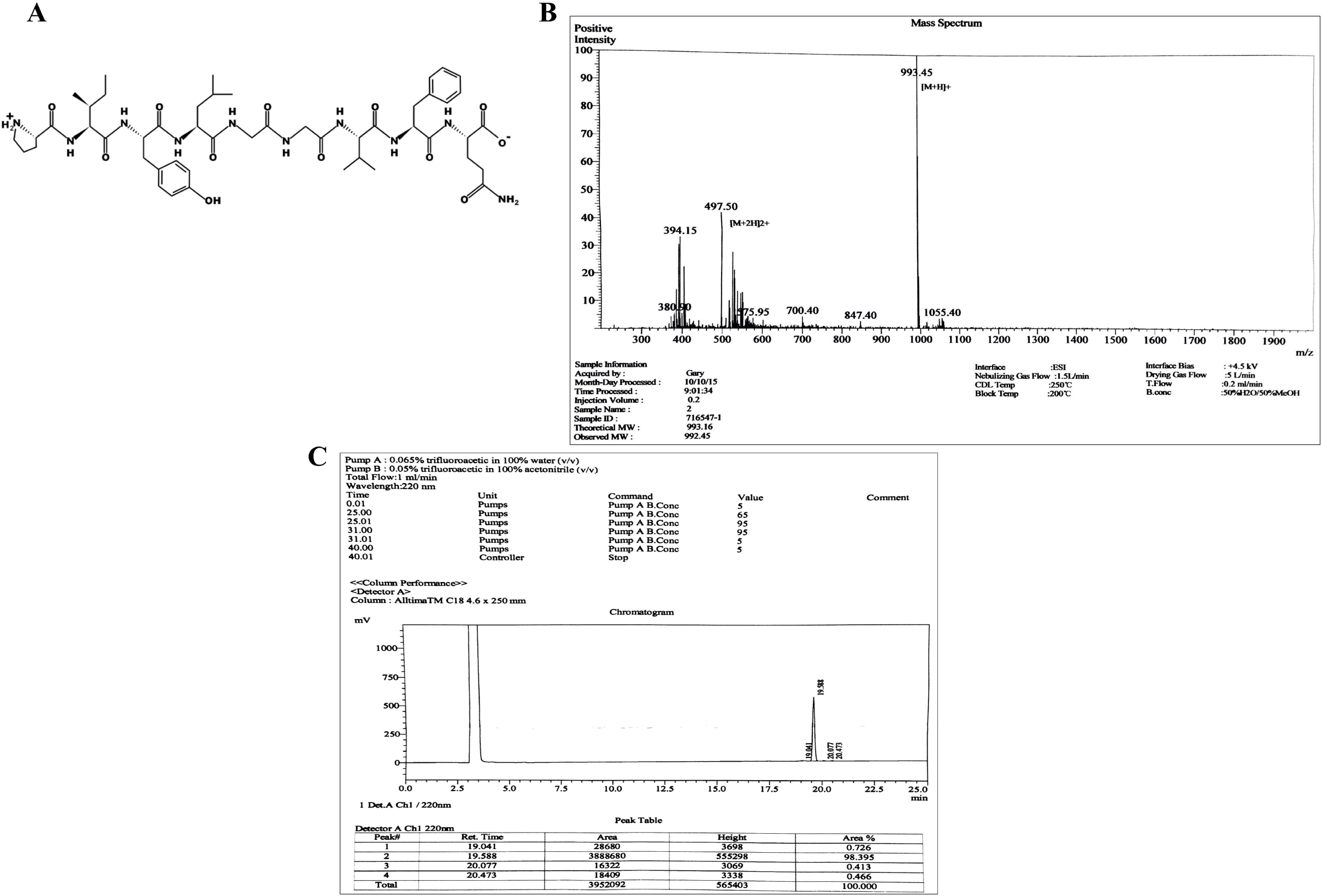
(A) Structure of the human anti-TNF-α peptide synthesized using solid-phase F-moc method. (B) Mass spectrometric profile of purified peptide, analysis was performed using MALDI TOF/TOF mass spectrometer. The molecular weight of the peptide is 993.45 as indicated. (C) Reverse-phase-HPLC profile of the anti-TNFα peptide after purification. The peak indicates the retention time.

**Fig S3.**
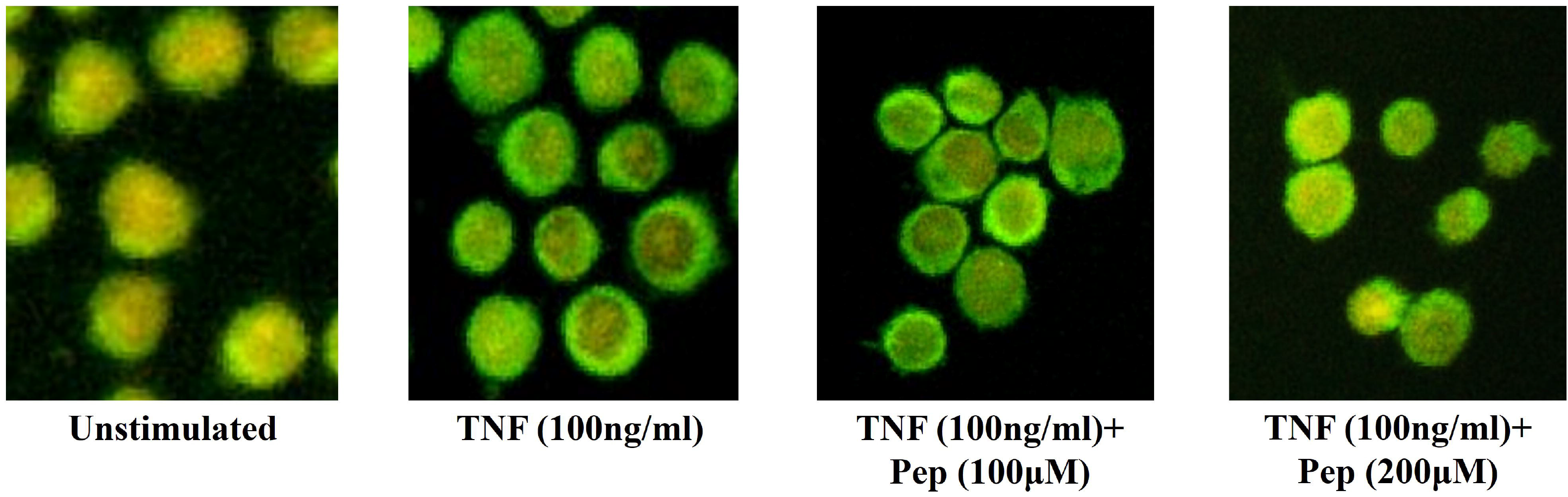
Inhibition of cell surface binding of TNF on U937 (human monocyte cell line) cells. The anti-TNFα peptide inhibits TNF-α binding to U937 human monocyte cells. The green signal is due to the TNF-α binding to the membrane receptors present on U937 cells (A). Cells not stimulated with TNFα, (B) Cells incubated with TNFα (100 ng/ml), and (C) Cells treated with the premixed TNFα and anti-TNFα peptide decrease the green signal (C).

**Figure S4.**
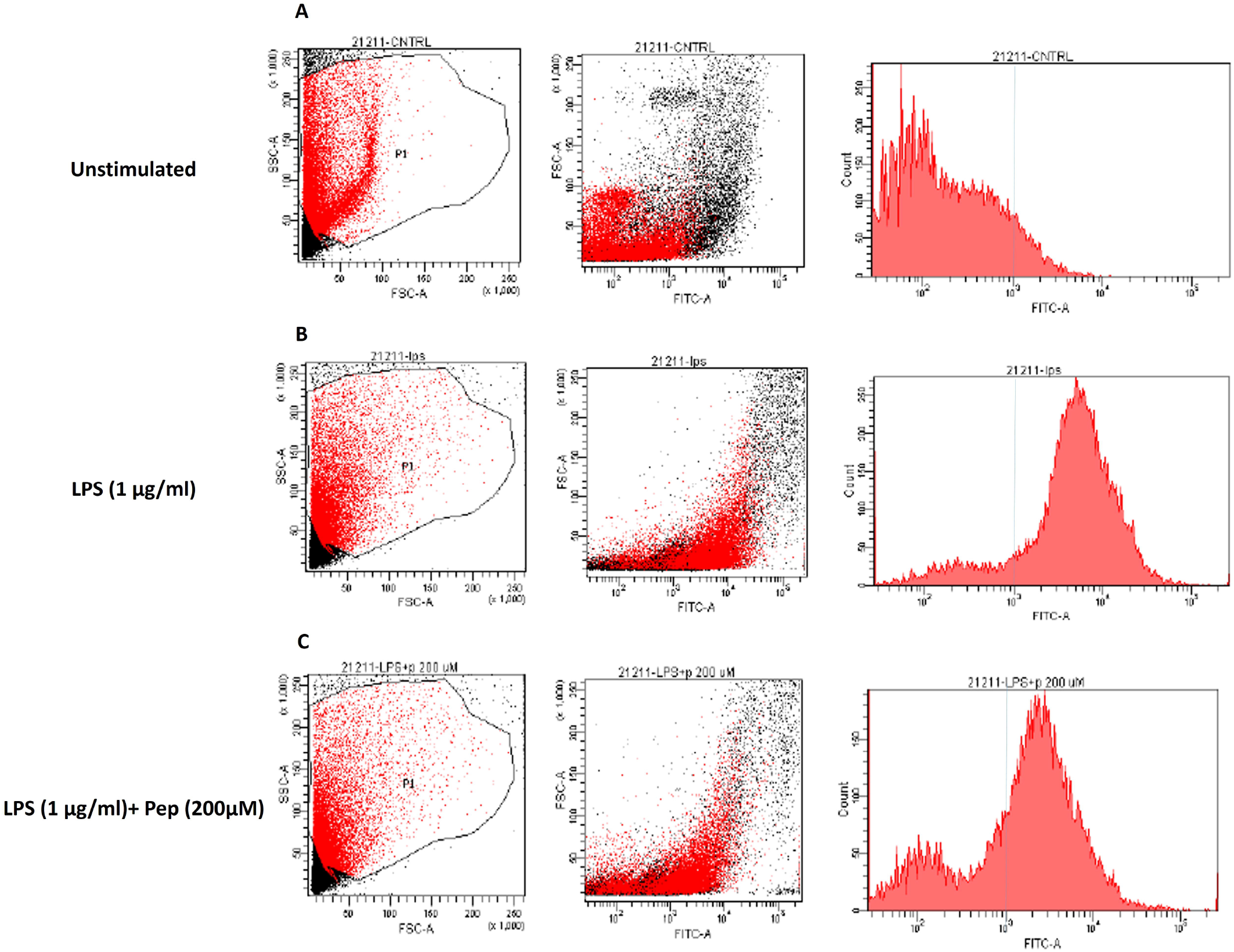
Histograms and dot plots from FACS enrichment analysis. (A) cells not stimulated with LPS, (B) cells stimulated with LPS (1μg/ml), and (C) LPS-stimulated (1 μg/ml) cells followed by treatment with peptide (200 μM) causing inhibition of TNF expression/binding on the cell surface.

