## Supplementary table for "Novel Peptide Inhibitor of Human Tumor Necrosis Factor-α has Antiarthritic Activity"

**Supplementary Tables**

**Table S1.** Peptides selected for *in-silico* synthesis and docking studies.

| **Peptide number** | **TNF strand** | **Stretch** | **Sequence** | **Length** |
| --- | --- | --- | --- | --- |
| 1 | e | 87-99 | YQTKVNLLSAIKS | 13 |
| 2 | e | 87-103 | YQTKVNLLSAIKSPCQR | 17 |
| 3 | e | 87-95 | YQTKVNLLS | 9 |
| 4 | e | 92-99 | NLLSAIKS | 8 |
| 5 | e | 92-103 | NLLSAIKSPCQR | 12 |
| 6 | f | 104-112 | KPWYEPIY | 8 |
| 7 | f | 104-119 | YLGGVFQ | 7 |
| 8 | f | 104-125 | KPWYEPIYLGGVFQ | 14 |
| **9** | **f** | **112-119** | **PIYLGGVFQ** | **9** |
| 10 | f | 119-125 | YEPIYLGGVFQ | 11 |
| 11 | f | 112-125 | SGQVYFGIIAL | 11 |
| 12 | f | 117-125 | ETPEGAEAK | 9 |
| 13 | F | 115-125 | ETPEGAEAKPWYEPIY | 16 |
| 14 | h | 147-157 | ETPEGAEAKPWYEPIYLGGVFQ | 22 |

**Table S2.**

|  | STRUCTURE: TNF α | | |  |  |  |  |  |  |  |  |
| --- | --- | --- | --- | --- | --- | --- | --- | --- | --- | --- | --- |
|  | Protein Residue Contacts | | | | | | A | B |  |  |  |
| Residue Number | | Residue Name | Chain ID | | Interface ASA | % Interface ASA | | | | Segment | H-Bonds |
| 53 | | GLU | A | | 27.56 | 2.17 | | | | 1 | . |
| 54 | | GLY | A | | 13.48 | 1.06 | | | | 1 | . |
| 55 | | LEU | A | | 75.89 | 5.99 | | | | 1 | . |
| 57 | | LEU | A | | 23.04 | 1.82 | | | | 1 | . |
| 72 | | THR | A | | 30.46 | 2.40 | | | | 2 | . |
| 73 | | HIS | A | | 50.07 | 3.95 | | | | 2 | 1 |
| 75 | | LEU | A | | 24.53 | 1.93 | | | | 2 | . |
| 82 | | ARG | A | | 16.64 | 1.31 | | | | 3 | 1 |
| 87 | | TYR | A | | 35.07 | 2.77 | | | | 3 | 1 |
| 91 | | VAL | A | | 22.51 | 1.78 | | | | 3 | . |
| 92 | | ASN | A | | 42.76 | 3.37 | | | | 3 | . |
| 93 | | LEU | A | | 30.39 | 2.40 | | | | 3 | . |
| 94 | | LEU | A | | 32.83 | 2.59 | | | | 3 | . |
| 95 | | SER | A | | 50.58 | 3.99 | | | | 3 | 1 |
| 96 | | ALA | A | | 28.89 | 2.28 | | | | 3 | . |
| 97 | | ILE | A | | 84.13 | 6.64 | | | | 3 | . |
| 98 | | LYS | A | | 48.49 | 3.82 | | | | 3 | . |
| 99 | | SER | A | | 40.75 | 3.21 | | | | 3 | . |
| 102 | | GLN | A | | 114.55 | 9.04 | | | | 3 | . |
| 103 | | ARG | A | | 109.38 | 8.63 | | | | 3 | 1 |
| 116 | | GLU | A | | 1.75 | 0.14 | | | | 4 | . |
| 118 | | ILE | A | | 3.33 | 0.26 | | | | 4 | . |
| 119 | | TYR | A | | 41.63 | 3.28 | | | | 4 | . |
| 120 | | LEU | A | | 15.57 | 1.23 | | | | 4 | . |
| 121 | | GLY | A | | 37.09 | 2.93 | | | | 4 | 1 |
| 122 | | GLY | A | | 17.26 | 1.36 | | | | 4 | . |
| 123 | | VAL | A | | 88.20 | 6.96 | | | | 4 | 1 |
| 124 | | PHE | A | | 48.35 | 3.81 | | | | 4 | . |
| 125 | | GLN | A | | 32.42 | 2.56 | | | | 4 | . |
| 157 | | LEU | A | | 80.19 | 6.32 | | | | 5 | . |

**Table S3.** PPI interaction server result, showing interface regions of A and C.

|  | STRUCTURE: TNF α | | |  |  |  |  |  |  |  |  |
| --- | --- | --- | --- | --- | --- | --- | --- | --- | --- | --- | --- |
|  | Protein Residue Contacts | | | | | | A | C |  |  |  |
| **Residue Number** | | **Residue Name** | **Chain ID** | | **Interface ASA** | **% Interface ASA** | | | | **Segment** | **H-Bonds** |
| 6 | | ARG | A | | 35.31 | 3.06 | | | | 1 | 1 |
| 8 | | PRO | A | | 76.31 | 6.61 | | | | 1 | . |
| 9 | | SER | A | | 17.99 | 1.56 | | | | 1 | . |
| 11 | | LYS | A | | 13.35 | 1.16 | | | | 1 | . |
| 13 | | VAL | A | | 20.70 | 1.79 | | | | 1 | . |
| 15 | | HIS | A | | 30.86 | 2.67 | | | | 1 | . |
| 33 | | ALA | A | | 3.25 | 0.28 | | | | 2 | . |
| 34 | | ASN | A | | 79.09 | 6.85 | | | | 2 | . |
| 36 | | LEU | A | | 30.62 | 2.65 | | | | 2 | . |
| 57 | | LEU | A | | 34.52 | 2.99 | | | | 3 | . |
| 59 | | TYR | A | | 45.00 | 3.90 | | | | 3 | . |
| 61 | | GLN | A | | 31.53 | 2.73 | | | | 3 | . |
| 63 | | LEU | A | | 22.56 | 1.96 | | | | 3 | . |
| 68 | | GLY | A | | 2.77 | 0.24 | | | | 3 | . |
| 69 | | CYS | A | | 4.42 | 0.38 | | | | 3 | . |
| 98 | | LYS | A | | 10.55 | 0.91 | | | | 4 | . |
| 101 | | CYS | A | | 12.08 | 1.05 | | | | 4 | . |
| 102 | | GLN | A | | 2.53 | 0.22 | | | | 4 | . |
| 103 | | ARG | A | | 6.16 | 0.53 | | | | 4 | . |
| 104 | | GLU | A | | 59.52 | 5.16 | | | | 4 | 1 |
| 109 | | ALA | A | | 22.06 | 1.91 | | | | 4 | . |
| 112 | | LYS | A | | 88.65 | 7.68 | | | | 4 | 1 |
| 113 | | PRO | A | | 35.96 | 3.12 | | | | 4 | 1 |
| 114 | | TRP | A | | 17.38 | 1.51 | | | | 4 | . |
| 115 | | TYR | A | | 88.13 | 7.64 | | | | 4 | . |
| 116 | | GLU | A | | 9.61 | 0.83 | | | | 4 | . |
| 117 | | PRO | A | | 55.60 | 4.82 | | | | 4 | . |
| 119 | | TYR | A | | 74.62 | 6.47 | | | | 4 | . |
| 143 | | LEU | A | | 2.08 | 0.18 | | | | 5 | . |
| 146 | | GLU | A | | 32.93 | 2.85 | | | | 5 | . |
| 148 | | GLY | A | | 69.01 | 5.98 | | | | 5 | 1 |
| 151 | | TYR | A | | 34.17 | 2.96 | | | | 5 | 1 |
| 155 | | ILE | A | | 59.69 | 5.17 | | | | 5 | . |
| 156 | | ALA | A | | 1.35 | 0.12 | | | | 5 | . |
| 157 | | LEU | A | | 23.52 | 2.04 | | | | 5 | . |

**Table S4.** PPI interaction server result, showing interface regions of B and A.

|  | STRUCTURE: TNF α | | |  |  |  |  |  |  |  |  |
| --- | --- | --- | --- | --- | --- | --- | --- | --- | --- | --- | --- |
|  | Protein Residue Contacts | | | | | | B | A |  |  |  |
| Residue Number | | Residue Name | Chain ID | | Interface ASA | % Interface ASA | | | | Segment | H-Bonds |
| 7 | | THR | B | | 18.71 | 1.53 | | | | 1 | . |
| 8 | | PRO | B | | 33.97 | 2.77 | | | | 1 | . |
| 9 | | SER | B | | 24.48 | 2.00 | | | | 1 | . |
| 11 | | LYS | B | | 28.00 | 2.29 | | | | 1 | . |
| 13 | | VAL | B | | 21.38 | 1.75 | | | | 1 | . |
| 14 | | ALA | B | | 1.43 | 0.12 | | | | 1 | . |
| 15 | | HIS | B | | 37.58 | 3.07 | | | | 1 | . |
| 33 | | ALA | B | | 34.98 | 2.86 | | | | 2 | 1 |
| 34 | | ASN | B | | 99.23 | 8.10 | | | | 2 | 1 |
| 36 | | LEU | B | | 69.64 | 5.69 | | | | 2 | . |
| 57 | | LEU | B | | 33.08 | 2.70 | | | | 3 | . |
| 59 | | TYR | B | | 36.02 | 2.94 | | | | 3 | 1 |
| 61 | | GLN | B | | 37.08 | 3.03 | | | | 3 | . |
| 63 | | LEU | B | | 22.23 | 1.81 | | | | 3 | . |
| 69 | | CYS | B | | 6.96 | 0.57 | | | | 4 | . |
| 98 | | LYS | B | | 15.01 | 1.23 | | | | 5 | . |
| 100 | | PRO | B | | 8.49 | 0.69 | | | | 5 | . |
| 104 | | GLU | B | | 66.44 | 5.42 | | | | 5 | 1 |
| 106 | | PRO | B | | 6.26 | 0.51 | | | | 5 | . |
| 109 | | ALA | B | | 20.10 | 1.64 | | | | 5 | . |
| 112 | | LYS | B | | 98.69 | 8.06 | | | | 5 | . |
| 113 | | PRO | B | | 31.26 | 2.55 | | | | 5 | 1 |
| 114 | | TRP | B | | 30.39 | 2.48 | | | | 5 | . |
| 115 | | TYR | B | | 89.00 | 7.27 | | | | 5 | . |
| 116 | | GLU | B | | 9.68 | 0.79 | | | | 5 | . |
| 117 | | PRO | B | | 46.18 | 3.77 | | | | 5 | . |
| 119 | | TYR | B | | 58.08 | 4.74 | | | | 5 | 1 |
| 143 | | LEU | B | | 3.23 | 0.26 | | | | 6 | . |
| 147 | | SER | B | | 51.17 | 4.18 | | | | 6 | . |
| 148 | | GLY | B | | 49.68 | 4.06 | | | | 6 | 1 |
| 149 | | GLN | B | | 37.37 | 3.05 | | | | 6 | . |
| 151 | | TYR | B | | 31.55 | 2.58 | | | | 6 | . |
| 155 | | ILE | B | | 44.19 | 3.61 | | | | 6 | . |
| 157 | | LEU | B | | 23.31 | 1.90 | | | | 6 | . |

**Table S5.** PPI interaction server result, showing interface regions of B and C.

|  | STRUCTURE: TNF α | | |  |  |  |  |  |  |  |
| --- | --- | --- | --- | --- | --- | --- | --- | --- | --- | --- |
|  | Protein Residue Contacts | | | B | C | |  |  |  |  |
| Residue Number | | Residue Name | Chain ID | | | Interface ASA | | % Interface ASA | Segment | H-Bonds |
| 53 | | GLU | B | | | 16.84 | | 1.48 | 1 | . |
| 54 | | GLY | B | | | 19.35 | | 1.70 | 1 | . |
| 55 | | LEU | B | | | 89.64 | | 7.89 | 1 | . |
| 57 | | LEU | B | | | 23.19 | | 2.04 | 1 | . |
| 72 | | THR | B | | | 5.72 | | 0.50 | 2 | . |
| 73 | | HIS | B | | | 46.51 | | 4.09 | 2 | 1 |
| 75 | | LEU | B | | | 27.88 | | 2.45 | 2 | . |
| 91 | | VAL | B | | | 13.43 | | 1.18 | 3 | . |
| 92 | | ASN | B | | | 21.90 | | 1.93 | 3 | 1 |
| 93 | | LEU | B | | | 28.47 | | 2.50 | 3 | . |
| 94 | | LEU | B | | | 30.68 | | 2.70 | 3 | . |
| 95 | | SER | B | | | 52.34 | | 4.60 | 3 | 1 |
| 96 | | ALA | B | | | 26.27 | | 2.31 | 3 | . |
| 97 | | ILE | B | | | 84.90 | | 7.47 | 3 | . |
| 98 | | LYS | B | | | 54.33 | | 4.78 | 3 | . |
| 99 | | SER | B | | | 30.08 | | 2.65 | 3 | 1 |
| 102 | | GLN | B | | | 97.33 | | 8.56 | 3 | . |
| 103 | | ARG | B | | | 37.25 | | 3.28 | 3 | . |
| 118 | | ILE | B | | | 4.27 | | 0.38 | 4 | . |
| 119 | | TYR | B | | | 46.89 | | 4.13 | 4 | . |
| 120 | | LEU | B | | | 11.82 | | 1.04 | 4 | . |
| 121 | | GLY | B | | | 41.90 | | 3.69 | 4 | 1 |
| 122 | | GLY | B | | | 16.29 | | 1.43 | 4 | . |
| 123 | | VAL | B | | | 88.35 | | 7.77 | 4 | 1 |
| 124 | | PHE | B | | | 47.52 | | 4.18 | 4 | . |
| 125 | | GLN | B | | | 70.73 | | 6.22 | 4 | 1 |
| 157 | | LEU | B | | | 102.71 | | 9.04 | 5 | 1 |

**Table S6.** PPI interaction server result, showing interface regions of C and A.

|  | STRUCTURE: TNF α | | |  |  |  |  |  |  |  |  |
| --- | --- | --- | --- | --- | --- | --- | --- | --- | --- | --- | --- |
|  | Protein Residue Contacts | | | | | | C | A |  |  |  |
| Residue Number | | Residue Name | Chain ID | | Interface ASA | % Interface ASA | | | | Segment | H-Bonds |
| 53 | | GLU | C | | 8.03 | 0.70 | | | | 1 | . |
| 54 | | GLY | C | | 15.95 | 1.38 | | | | 1 | . |
| 55 | | LEU | C | | 81.98 | 7.11 | | | | 1 | . |
| 57 | | LEU | C | | 25.12 | 2.18 | | | | 1 | . |
| 72 | | THR | C | | 1.10 | 0.10 | | | | 2 | . |
| 73 | | HIS | C | | 55.02 | 4.77 | | | | 2 | 1 |
| 75 | | LEU | C | | 18.57 | 1.61 | | | | 2 | . |
| 82 | | ARG | C | | 4.44 | 0.39 | | | | 3 | . |
| 92 | | ASN | C | | 13.93 | 1.21 | | | | 4 | . |
| 93 | | LEU | C | | 34.03 | 2.95 | | | | 4 | . |
| 94 | | LEU | C | | 22.53 | 1.96 | | | | 4 | . |
| 95 | | SER | C | | 40.20 | 3.49 | | | | 4 | 1 |
| 96 | | ALA | C | | 32.83 | 2.85 | | | | 4 | . |
| 97 | | ILE | C | | 104.55 | 9.07 | | | | 4 | . |
| 98 | | LYS | C | | 54.26 | 4.71 | | | | 4 | . |
| 99 | | SER | C | | 32.90 | 2.86 | | | | 4 | . |
| 101 | | CYS | C | | 4.70 | 0.41 | | | | 4 | 1 |
| 102 | | GLN | C | | 119.52 | 10.37 | | | | 4 | . |
| 103 | | ARG | C | | 64.96 | 5.64 | | | | 4 | 1 |
| 118 | | ILE | C | | 4.74 | 0.41 | | | | 5 | . |
| 119 | | TYR | C | | 38.60 | 3.35 | | | | 5 | . |
| 120 | | LEU | C | | 14.30 | 1.24 | | | | 5 | . |
| 121 | | GLY | C | | 41.25 | 3.58 | | | | 5 | 1 |
| 122 | | GLY | C | | 14.11 | 1.22 | | | | 5 | . |
| 123 | | VAL | C | | 83.02 | 7.20 | | | | 5 | . |
| 124 | | PHE | C | | 48.65 | 4.22 | | | | 5 | . |
| 125 | | GLN | C | | 76.40 | 6.63 | | | | 5 | 1 |
| 127 | | GLU | C | | 4.49 | 0.39 | | | | 5 | . |
| 157 | | LEU | C | | 92.21 | 8.00 | | | | 6 | . |

**Table S7.** PPI interaction server result, showing interface regions of C and B.

|  | STRUCTURE: TNF α | | |  |  |  |  |  |  |  |  |
| --- | --- | --- | --- | --- | --- | --- | --- | --- | --- | --- | --- |
|  | Protein Residue Contacts | | | | | | C | B |  |  |  |
| Residue Number | | Residue Name | Chain ID | | Interface ASA | % Interface ASA | | | | Segment | H-Bonds |
| 6 | | ARG | C | | 31.49 | 2.84 | | | | 1 | 1 |
| 8 | | PRO | C | | 57.13 | 5.15 | | | | 1 | . |
| 9 | | SER | C | | 48.06 | 4.34 | | | | 1 | 1 |
| 11 | | LYS | C | | 22.99 | 2.07 | | | | 1 | . |
| 13 | | VAL | C | | 21.55 | 1.94 | | | | 1 | . |
| 15 | | HIS | C | | 34.86 | 3.14 | | | | 1 | . |
| 33 | | ALA | C | | 15.73 | 1.42 | | | | 2 | . |
| 34 | | ASN | C | | 71.89 | 6.48 | | | | 2 | . |
| 36 | | LEU | C | | 23.39 | 2.11 | | | | 2 | . |
| 57 | | LEU | C | | 29.22 | 2.64 | | | | 3 | . |
| 59 | | TYR | C | | 43.44 | 3.92 | | | | 3 | 1 |
| 61 | | GLN | C | | 33.68 | 3.04 | | | | 3 | . |
| 63 | | LEU | C | | 17.02 | 1.54 | | | | 3 | . |
| 69 | | CYS | C | | 5.30 | 0.48 | | | | 4 | . |
| 98 | | LYS | C | | 13.49 | 1.22 | | | | 5 | . |
| 100 | | PRO | C | | 10.23 | 0.92 | | | | 5 | . |
| 104 | | GLU | C | | 47.26 | 4.26 | | | | 5 | . |
| 109 | | ALA | C | | 3.29 | 0.30 | | | | 5 | . |
| 112 | | LYS | C | | 61.63 | 5.56 | | | | 5 | . |
| 113 | | PRO | C | | 34.51 | 3.11 | | | | 5 | 2 |
| 114 | | TRP | C | | 26.87 | 2.42 | | | | 5 | . |
| 115 | | TYR | C | | 87.04 | 7.85 | | | | 5 | . |
| 116 | | GLU | C | | 7.93 | 0.72 | | | | 5 | . |
| 117 | | PRO | C | | 56.99 | 5.14 | | | | 5 | . |
| 119 | | TYR | C | | 61.97 | 5.59 | | | | 5 | 1 |
| 143 | | LEU | C | | 3.22 | 0.29 | | | | 6 | . |
| 147 | | SER | C | | 34.74 | 3.13 | | | | 6 | 1 |
| 148 | | GLY | C | | 52.27 | 4.71 | | | | 6 | 1 |
| 149 | | GLN | C | | 40.36 | 3.64 | | | | 6 | . |
| 151 | | TYR | C | | 33.12 | 2.99 | | | | 6 | . |
| 155 | | ILE | C | | 52.22 | 4.71 | | | | 6 | . |
| 157 | | LEU | C | | 25.75 | 2.32 | | | | 6 | . |
