## Supplementary material for "Novel Peptide Inhibitor of Human Tumor Necrosis Factor-α has Antiarthritic Activity": Table 1

**Table 1.** The binding energy between TNFα and ligands (Std and Peptide) interactions, as derived from the Molecular Dynamics studies.

| **Energy Component (in kcal/mol)** | **STD** | **Peptide** |
| --- | --- | --- |
| **E_VDWAALS_** | -47.66 | -68.88 |
| **E_EL_** | -5.60 | -86.73 |
| **E_GB_** | 15.80 | 127.96 |
| **E_SURF_** | -5.85 | -10.57 |
| **ΔG_gas_** | -53.25 | 155.60 |
| **ΔG_solv_** | 9.95 | 117.39 |
| **ΔG_bind_** | **-43.31** | **-38.21** |
